# MAVERICC: Efficient marker-free rescue of vaccinia virus recombinants by *in vitro* CRISPR-Cas9 engineering

**DOI:** 10.1101/2021.01.05.425447

**Authors:** Ethan Laudermilch, Kartik Chandran

**Affiliations:** Department of Microbiology and Immunology, Albert Einstein College of Medicine, Bronx, NY 10461

## Abstract

Vaccinia virus (VACV)-based vectors are in extensive use as vaccines and cancer immunotherapies. VACV engineering has traditionally relied on homologous recombination between a parental viral genome and a transgene-bearing transfer plasmid, a highly inefficient process that necessitates the use of a selection or screening marker to isolate recombinants. Recent extensions of this approach have sought to enhance the recovery of transgene-bearing viruses through the use of CRISPR-Cas9 engineering to cleave the viral genome in infected cells. However, these methods do not completely eliminate the generation of WT viral progeny and thus continue to require multiple rounds of viral propagation and plaque purification. Here, we describe MAVERICC (<u>ma</u>rker-free <u>v</u>accinia virus <u>e</u>ngineering of recombinants through *in vitro* <u>C</u>RISPR/Cas9 <u>c</u>leavage), a new strategy to engineer recombinant VACVs in a manner that overcomes current limitations. MAVERICC also leverages the CRISPR/Cas9 system but requires no markers and yields essentially pure preparations of the desired recombinants in a single step. We used this approach to rapidly introduce point mutations, insertions, and deletions at multiple locations in the VACV genome, both singly and in combination. The efficiency and versatility of MAVERICC make it an ideal choice for generating mutants and mutant libraries at arbitrarily selected locations in the viral genome to build complex VACV vectors, effect vector improvements, and facilitate the study of poxvirus biology.

**Graphical Abstract:** 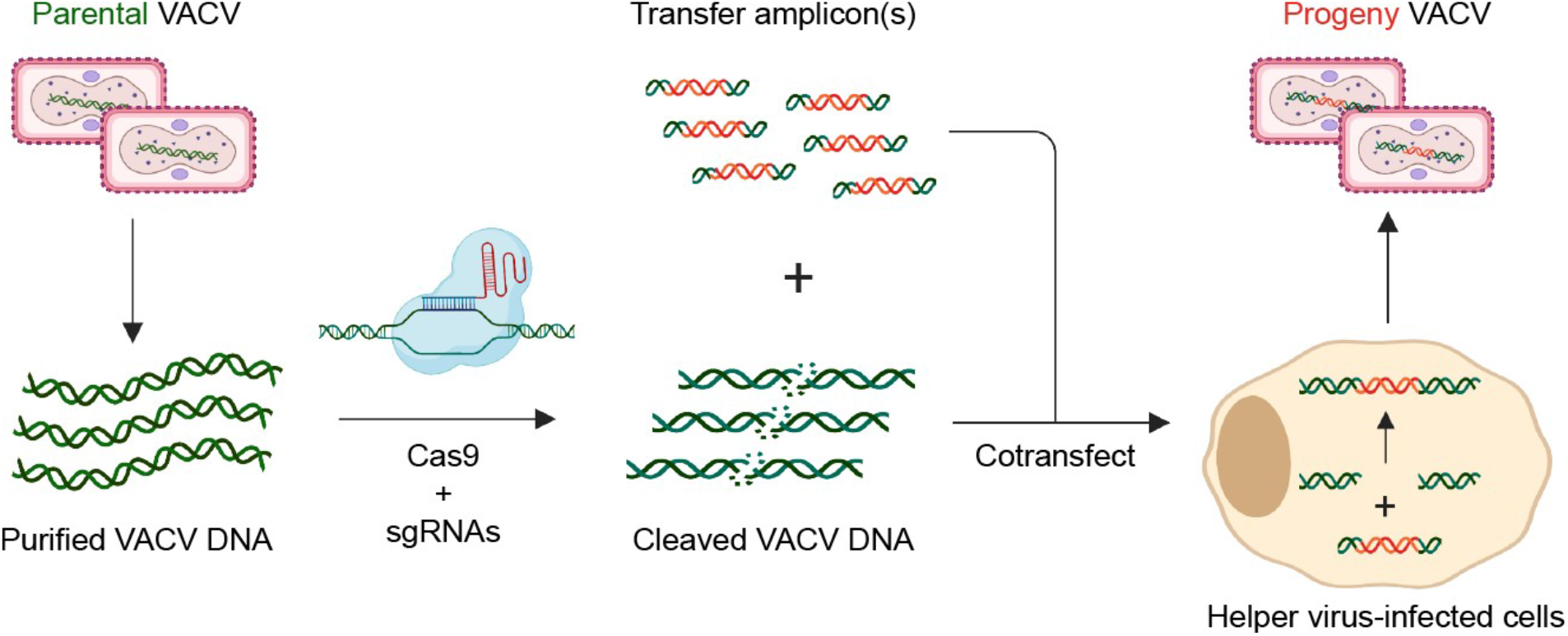

**Overview of MAVERICC.** Conceptual overview of the approach outlined in this manuscript. To make VACV recombinants, the parental virus is first purified and vDNA is isolated with phenol:chloroform extraction. This purified vDNA is then treated with Cas9 enzyme and sgRNAs that are directed to a specific locus in the VACV genome. The cleaved vDNA is then transfected into FWPV-infected BSC-40 cells along with a transfer amplicon containing an insertion or mutation of interest flanked by homologous sequences. Recombination is allowed to occur for 5-7 days, during which time the cleaved vDNA is healed by the transfer amplicon, thus editing the VACV genome, and packaged into infectious viral particles. Individual plaques are grown up and rVACVs are isolated after a single round of plaque purification. Image was created with Biorender.com.

## Introduction

Vaccinia virus (VACV), the prototypical orthopoxvirus, has a large linear DNA genome of nearly 200 kb and encodes approximately 250 genes. Like other poxviruses, VACV replicates entirely in the cytoplasm and encodes its own machinery for genome transcription and replication. Historically, VACV is perhaps most well known as a successful vaccine in the global campaign to eradicate smallpox, which was finally achieved in the 1970s. Although routine smallpox vaccination has ceased, VACV has continued to enjoy widespread popularity as an expression vector for vaccination [1] and cancer immunotherapy [2,3]. At least one VACV-vectored vaccine is approved for use in animals and multiple others are currently under development for use in animals and humans [4,5]. Features that make VACV a popular choice for such applications include its broad host range (including most human cells) [6], capacity to harbor up to ∼25 kb in foreign genetic material [7], thermal stability [8], and ease of propagation [9]. Moreover, as a vaccine and immunotherapy vector, VACV induces strong and durable cellular and humoral immune responses to heterologous antigens [1].

Despite their widespread use, current methods for generating recombinant vaccinia viruses (rVACVs) are limited in efficiency and/or versatility. Several afford the engineering of a wide variety of rVACVs but can be slow and cumbersome. These include the traditional approach [9], in which cells are infected with WT virus and then transfected with a transfer plasmid that contains homology arms flanking a foreign gene or mutated VACV gene and a selectable marker. Homologous recombination between viral genomic DNA (vDNA) and the transfer plasmid, a rare event, yields about 1 recombinant clone for every 1,000 WT progeny virions. Recombinant clones are enriched through marker selection or screening and multiple rounds of plaque purification in a process that typically takes weeks.

Recently, CRISPR/Cas9 engineering [10] has been employed in attempts to streamline the rVACV rescue process [11–15]. Yuan and co-workers [11] introduced *Streptococcus pyogenes* Cas9 and vDNA-specific single-guide RNAs (sgRNAs) into cells to cleave the vDNA from infecting VACVs and enhance its homologous recombination with a transfected transfer plasmid. This approach marginally increased the efficiency of rVACV formation over the traditional method but still required the insertion, and possible subsequent removal, of a marker gene. An elaboration of this strategy used intracellular vDNA cleavage by Cas9 to selectively inhibit the replication of parental genomes, thereby affording the marker-free recovery of rVACVs [13]. However, multiple rounds of selection were still required to obtain single-site recombinants at a high frequency, and significant levels of WT contamination persisted, rendering this approach unsuitable for the generation of rVACV libraries.

Other rVACV rescue approaches have overcome the efficiency problem described above but at the expense of versatility. In one strategy, termed trimolecular recombination [16,17], purified vDNA is cleaved with restriction enzymes and introduced into cells along with a transfer plasmid or linear amplicon that serves as a repair template. A replication-defective helper poxvirus provides the trans-acting factors that heal the cleaved VACV DNA by effecting a three-way recombination reaction. Trimolecular recombination is highly efficient and generates essentially no WT background, making it suitable for recovery of both single recombinants and viral libraries. However, its use is limited by the availability of unique restriction sites, which are rare in the nearly 200-kb VACV genome and typically must first be engineered into a specific locus. Thus, this approach is most useful for insertion of foreign genes. A second recently developed approach, EPICC (efficient purification by parental inducer constraint), uses a replication-inducible rVACV as a parental virus for homologous recombination [18]. However, like trimolecular recombination, this method’s utility is largely limited to ‘special’ modified loci in the vDNA that can serve as landing pads for heterologous DNA sequences.

Here, we describe a new strategy to generate rVACVs that overcomes the limitations of existing platforms by combining high efficiency and versatility. MAVERICC (<u>ma</u>rker-free <u>v</u>accinia virus <u>e</u>ngineering of recombinants through *in vitro* <u>C</u>RISPR/Cas9 <u>c</u>leavage) produced recombinants bearing the desired point mutations, insertions, or deletions at different locations in the VACV genome with >90% recovery in less than a week, despite requiring no markers, special cell lines, engineered parental viruses, serial passages, or selection steps. Moreover, its efficiency afforded the simultaneous engineering of two distinct genomic loci with ∼70% recovery. We anticipate that MAVERICC will facilitate the rapid, marker-free construction of complex rVACV vectors as well as viral libraries to iteratively improve these vectors and investigate the basic biology of poxviruses.

## Results

### Site-specific, sgRNA-directed Cas9 cleavage of VACV genomic DNA *in vitro*

As an initial step towards establishing MAVERICC, we tested if VACV genomic DNA (vDNA) could be efficiently targeted with CRISPR/Cas9 *in vitro*. We first designed single-guide RNAs (sgRNAs) to excise the *mCherry* reporter gene from the thymidine kinase (*tk*) locus of a previously engineered parental virus (Figure 1(A)). To minimize off-target cleavage, sgRNA sequences were screened with CRISPOR [19] to identify those that contained 6 or more mismatches to sequences elsewhere in the VACV genome. vDNA isolated from the parental virus [20,21] was incubated with purified Cas9 and the sgRNAs, and then digested into 16 previously defined fragments with the *HindIII* restriction enzyme [20]. The cleavage products were resolved by agarose gel electrophoresis to visualize the 5,800 bp *HindIII* J fragment, which contains the *tk* locus, and therefore the *mCherry* sequence. Cas9+sgRNA treatment resulted in essentially complete loss of this fragment, indicating that the vDNA was specifically and efficiently cleaved (Figure 1(B)).

**Figure 1.**
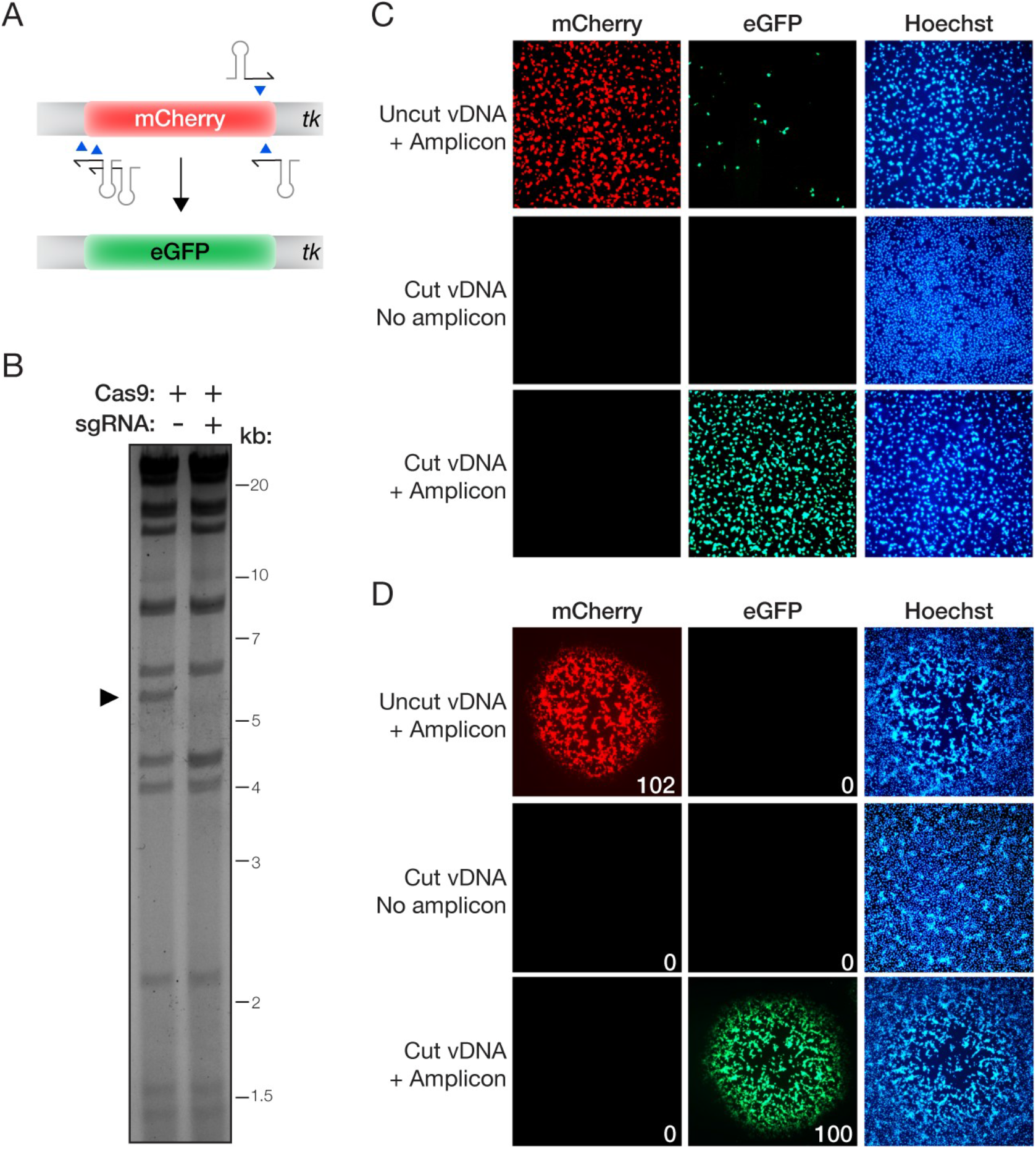
Efficient exchange of fluorescent reporters with MAVERICC. (A) Overview of the strategy to exchange an eGFP reporter for an mCherry reporter currently in the tk locus. The parental vDNA was cut with Cas9 directed by four sgRNAs (stem-loop and red arrowhead) as shown. (B) vDNA (incubated with Cas9 without or with sgRNAs) was fragmented by HindIII digestion, resolved by agarose gel electrophoresis, and visualized by ethidium bromide staining. Black arrowhead, DNA fragment containing the mCherry reporter gene. (C) Fresh cells were exposed to viral rescue samples and examined for mCherry or eGFP reporter expression by fluorescence microscopy at 24 h post-infection. (D) Viral rescue samples were subjected to plaque assay and individual viral plaques were randomly selected and enumerated as mCherry-positive or eGFP-positive (total red or green plaques shown in the lower right corner of individual plaque images). No plaques were observed in rescue samples that received cleaved vDNA but no transfer amplicon (middle row).

### Generation of a fluorescent marker-swapped rVACV

We next investigated if this Cas9-cleaved vDNA could serve as a substrate for trimolecular recombination in cells to generate a rVACV in which the *mCherry* reporter gene was replaced by *eGFP*. Accordingly, we prepared a transfer amplicon bearing the *eGFP* sequence flanked by ∼300-bp homology arms specific for the *tk* locus to use as a repair template. Cells were infected with a replication-defective helper poxvirus (fowlpox virus; FWPV), transfected with cleaved vDNA and the transfer amplicon, and incubated for 6 days to allow recombination to occur. Cells and cell supernatants were then harvested and exposed to fresh cells for 24 h to detect and amplify any infectious viral progeny (Figure 1(C)). As expected, the cells co-transfected with uncleaved vDNA and the transfer amplicon generated mostly mCherry-expressing virions, although a few cells expressing eGFP were also detected. By contrast, cells receiving Cas9-cleaved vDNA and the transfer amplicon generated only eGFP-expressing virions. Introduction of cleaved vDNA into cells without the transfer amplicon produced little or no infectious virus, attesting to the high efficiency of *in vitro* Cas9 cleavage. To confirm that the fluorescent reporter expression observed above arose from infection by replication-competent viral clones, we grew up individual plaques for 3 days under a 0.5% methylcellulose overlay. Concordantly, all of the ∼100 plaques we counted from the sample co-transfected with uncleaved vDNA and the eGFP transfer amplicon produced only mCherry-positive plaques, whereas those from the sample co-transfected with cleaved vDNA and the eGFP transfer amplicon were all eGFP-positive (Figure 1(D)). No viral plaques were observed from the sample transfected with cleaved vDNA alone. Thus, sgRNA-programmed Cas9 cleavage of vDNA *in vitro* affords the recovery of a desired rVACV recombinant with high efficiency and little or no WT contamination.

### Marker-free engineering of VACV gene products involved in the biogenesis of extracellular enveloped virions

The preceding findings indicated that our approach could be amenable to engineering the VACV genome in an altogether marker-free manner. To test this hypothesis, we introduced mutations in two VACV genes previously shown to enhance the secretion of extracellular enveloped virions (EEVs) from infected cells [22–24]. Specifically, we designed constructs to truncate the *A33R* gene at the C–terminus (ΔCT) and mutate the *A34R* gene at amino acid residue 151 (K151E; AAA codon to GAA). Each locus was targeted with two different sgRNAs predicted to specifically recognize sequences near the mutation site (Figure 2(A)). We incubated purified vDNA with Cas9 and these sgRNAs as above, and then fragmented the vDNA with the *XhoI* restriction enzyme for analysis. *A33R* and *A34R* both reside in a 10.9-kb *XhoI* fragment, and its sgRNA-programmed cleavage by Cas9 yielded products of 8.2 kb and 2.7 kb (*A33R*) or 8.9 kb and 2.0 kb (*A34R*), exactly as predicted (Figure 2(B)). We next co-transfected FWPV-infected BSC-40 cells with the Cas9-cleaved vDNA and a transfer amplicon harboring the desired mutation and flanked by 500-bp homology arms. After 5 days, we observed robust viral gene expression in wells that received the cleaved vDNA and the PCR amplicon but not in wells that received only cleaved vDNA (data not shown), indicating that rVACV rescue was successful.

**Figure 2.**
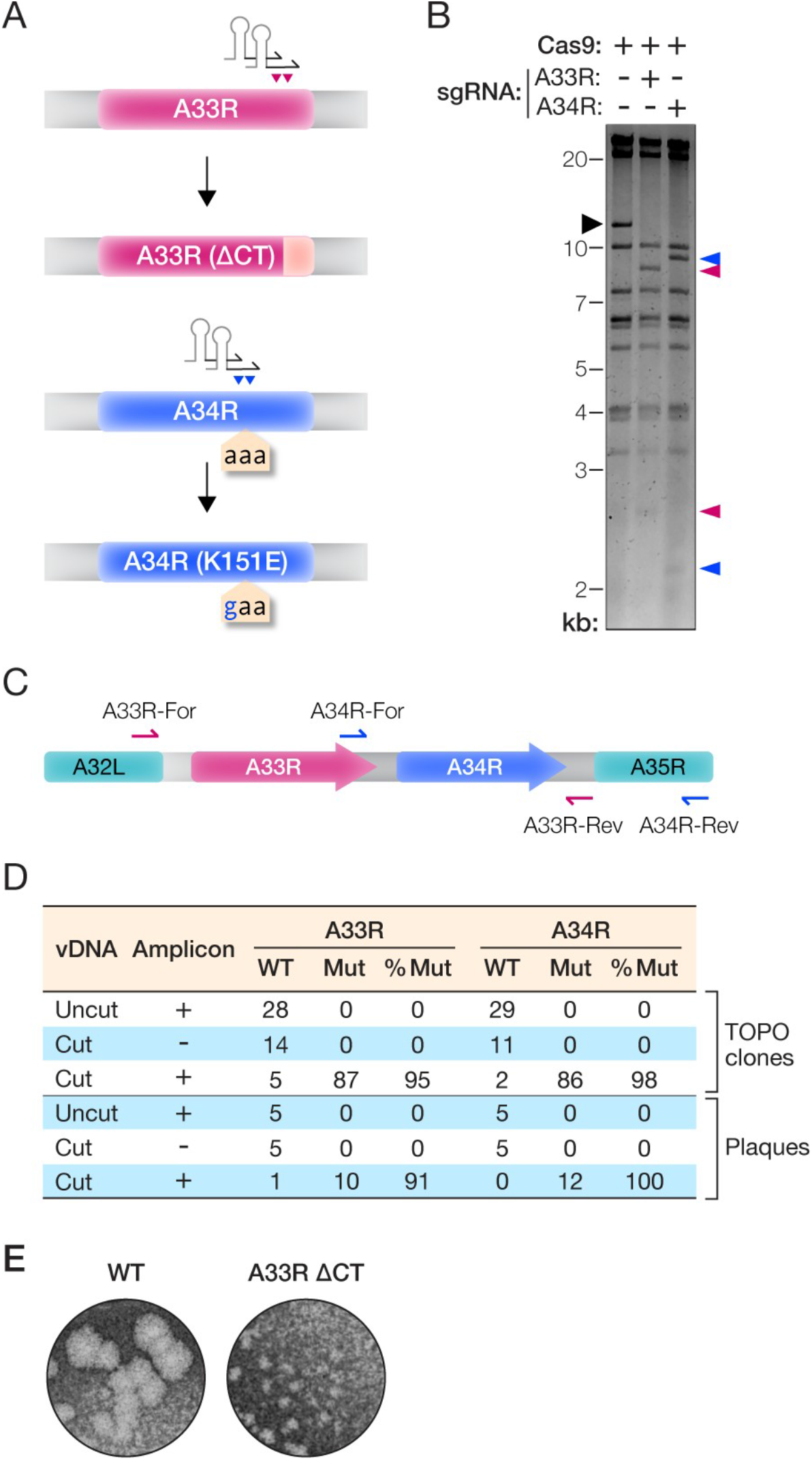
Mutation of A33R and A34R genes without a selectable marker. (A) Overview of the strategy to introduce mutations into the A33R or A34R genes. (B) vDNA (incubated with Cas9 without or with sgRNAs) was fragmented by XhoI digestion, resolved by agarose gel electrophoresis, and visualized by ethidium bromide staining. Black arrowhead, vDNA fragment containing the A33R and A34R genes. Pink and blue arrowheads, new vDNA fragments observed upon cleavage with Cas9 and sgRNAs specific for A33R or A34R, respectively. (C) Overview of the PCR strategy to amplify the A33R or A34R loci from the mixed population of viral genomes after rescue of transfected vDNA by MAVERICC. Primers annealing outside the transfer amplicons were used (pink or blue arrows) (D) PCR amplicons from panel C were sequenced to determine the efficiency of rVACV rescue. The number of plasmids or PCR amplicons showing WT or mutant sequences and the corresponding percentage of mutant sequences are shown. (E) WT and A33R(ΔCT) mutant plaques were visualized by staining with crystal violet after 4 days of growth under a 0.5% methylcellulose overlay.

We used two approaches to estimate the efficiency of trimolecular recombination and incorporation of the desired mutations into rVACVs. First, we purified vDNA from cell suspensions harvested at 5 days post-rescue and subjected them to PCR with primers designed to selectively amplify the genomic *A33R* or *A34R* loci (Figure 2(C)). To obtain a representative sample, cycling conditions were optimized to keep product amplification in the linear range and three independent reactions were pooled prior to analysis (Figure S1). Individual amplicons were isolated by TOPO cloning and sequenced. Only WT *A33R* and *A34R* sequences were obtained from cells that received either uncleaved vDNA with transfer amplicon or cleaved vDNA without transfer amplicon. By contrast, we observed the A33R(ΔCT) and A34R(K151E) mutations in >95% of TOPO clones from cells that received both cleaved vDNA and the respective transfer amplicon, indicating that homologous recombination had occurred with high efficiency (Figure 2(D)).

In the second approach, we attempted to isolate mutant rVACV clones without employing a marker for screening or selection. Accordingly, we subjected the cell suspensions harvested at 5 days post-rescue to a plaque assay, picked and amplified individual plaques at random, and sequenced the *A33R* and *A34R* genomic loci in vDNA prepared from these viral stocks. Consistent with our findings above, we obtained only WT rVACV clones from cells that received either uncleaved vDNA with transfer amplicon or cleaved vDNA without transfer amplicon, whereas >90% of clones from cells that received both cleaved vDNA and transfer amplicon bore the desired genomic mutation (Figure 2(D)). As shown previously [23,24], rVACVs bearing A33R(ΔCT) displayed a small-plaque phenotype (Figure 2(E)), providing independent confirmation that we had successfully engineered them to encode and express this mutant protein. Therefore, MAVERICC affords the highly efficient, marker-free construction of rVACVs in a single step.

### Generating rVACVs bearing epitope-tagged proteins in the entry fusion complex (EFC)

We postulated that MAVERICC could be generalized to other regions of the VACV genome. To test this hypothesis, we attempted to introduce a triple FLAG epitope tag sequence (3X-FLAG) at the C-terminus of H2R and L5R (Figure 3(A)), two EFC proteins that are each essential for VACV entry [25] into cells. Applying the MAVERICC protocol to an eGFP-expressing parental virus, we observed Cas9 cleavage of the expected DNA fragments with sgRNAs designed to target the *H2R* or *L5R* loci (Figure 3(B)). After rescue, we observed robust viral gene expression in the cells from the H2R-3X FLAG rescue experiment but no evidence of viral replication in the L5R-3X FLAG cells (data not shown). To investigate our failure to recover virus containing epitope-tagged L5R, we assessed recombination efficiency by PCR-amplifying the *H2R* and *L5R* genomic loci in vDNA isolated from the cell suspensions at 5 days post-rescue. For both rescue experiments, cells that received cleaved vDNA and transfer amplicon yielded a larger PCR product than those that received only cleaved vDNA (Figure 3(C)). Sanger sequencing of these products confirmed that the FLAG tag-encoding sequence was present as in-frame fusions to *H2R* and *L5R* in the larger PCR amplicons, despite the lack of detectable viral replication in the *L5R* sample (data not shown). Concordantly, we obtained rVACV clones encoding H2R-3X FLAG from 11 of 12 isolated plaques. The H2R-3X FLAG protein migrated at the expected molecular weight and localized to sites of viral assembly (Fig 3(D–E)). Unexpectedly, however, we observed no plaque formation in samples from the L5R-3X FLAG rescue experiment. We conclude that insertion of the 3X-FLAG sequence into the essential *L5R* gene was successful but lethal. Our attemptto engineer *L5R* in this manner also did not generate any WT clones, demonstrating once again the high efficiency of rVACV engineering afforded by trimolecular recombination in general and MAVERICC in particular.

**Figure 3.**
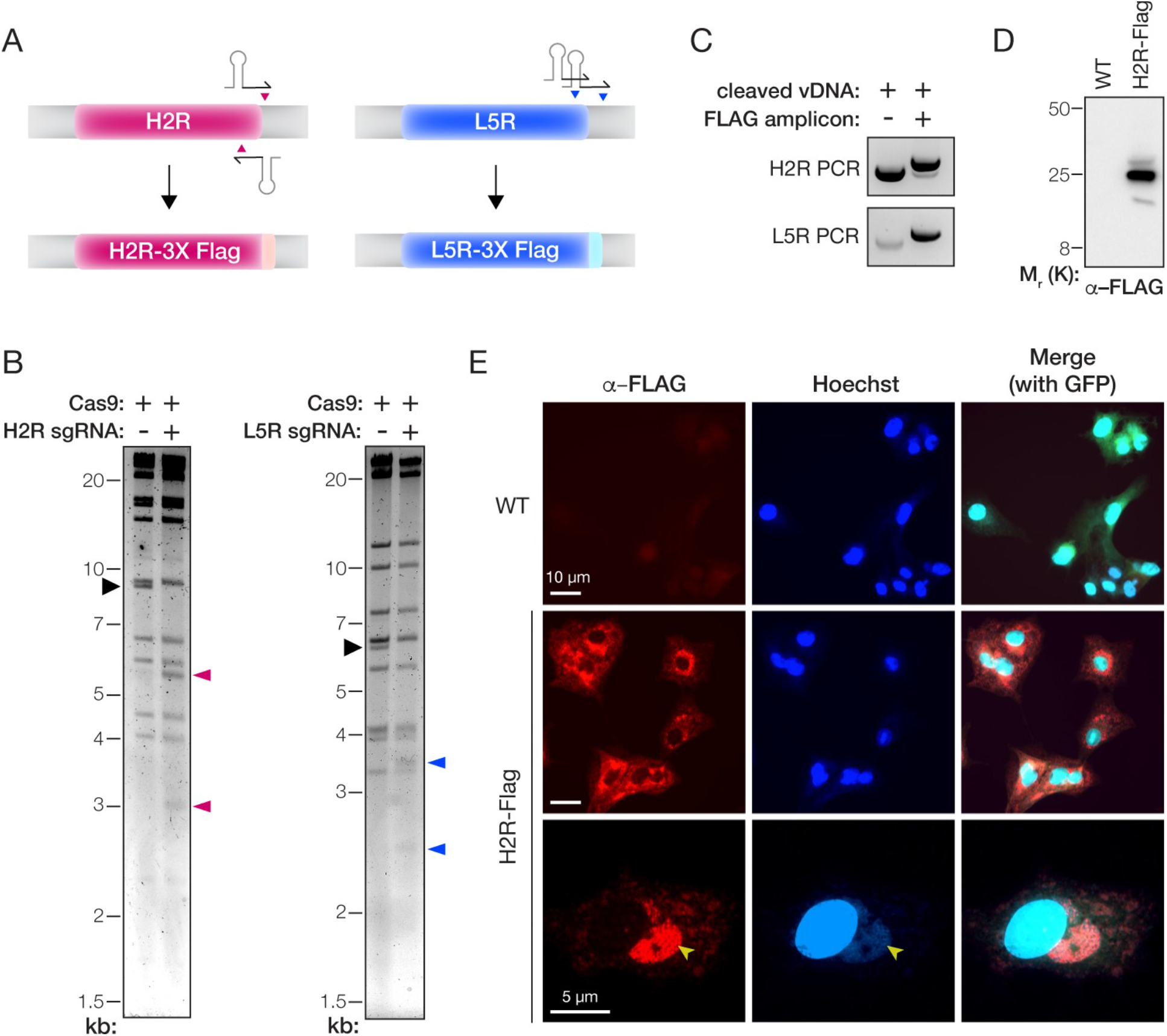
Epitope-tagging of genes encoding components of the entry fusion complex. (A) Depiction of the strategy to fuse a 3X-FLAG tag to the C–terminus of the H2R and L5R proteins. (B) vDNA (incubated with Cas9 without or with sgRNAs) was fragmented by digestion with HindIII (H2R) or XhoI (L5R) digestion, resolved by agarose gel electrophoresis, and visualized by ethidium bromide staining. Black arrowhead, fragment harboring the H2R or L5R gene. Pink or blue arrowheads: vDNA fragments observed upon cleavage with Cas9 and sgRNAs specific for H2R or L5R, respectively. (C) PCR amplicons produced by amplifying the mixed population of viral genomes with gene-specific primers after rescue of transfected vDNA (without or with transfer amplicon) by MAVERICC. (D) Extracts of cells infected with WT or H2R-FLAG-tagged viruses were resolved on a polyacrylamide gel and FLAG-tagged proteins were visualized by immunoblotting. (E) Images of BSC-40 cells infected with WT or H2R-FLAG-tagged viruses and immunostained with α–FLAG antibody. Cells were counterstained with Hoechst 33342 to visualize nuclei and cytoplasmic sites of viral assembly (yellow arrowheads). eGFP expressed from the tk locus of both viruses is used as a marker for viral infection in the merge panels at the right.

### Applying MAVERICC to two genes simultaneously

Finally, given the high efficiency of making a single genome edit, we sought to introduce multiple concurrent changes to the VACV genome. To this end, we incubated vDNA with sgRNA-loaded Cas9 specific for both *A33R* and *H2R* before rescuing these constructs with A33(ΔCT) and H2R-3X FLAG transfer amplicons, as described above. We then selected and amplified individual plaques at random. Sequencing of the *A33R* and *H2R* genomic loci in vDNA prepared from these viral stocks revealed that ∼80% and ∼90% of the plaques contained the desired *A33R* and *H2R* changes, respectively. Importantly, ∼70% of the analyzed plaques contained both mutations (Figure 4). Therefore, MAVERICC affords the rapid recovery of rVACVs bearing multiple engineered genomic loci in a single step without the need for any selection or screening markers.

**Figure 4.**
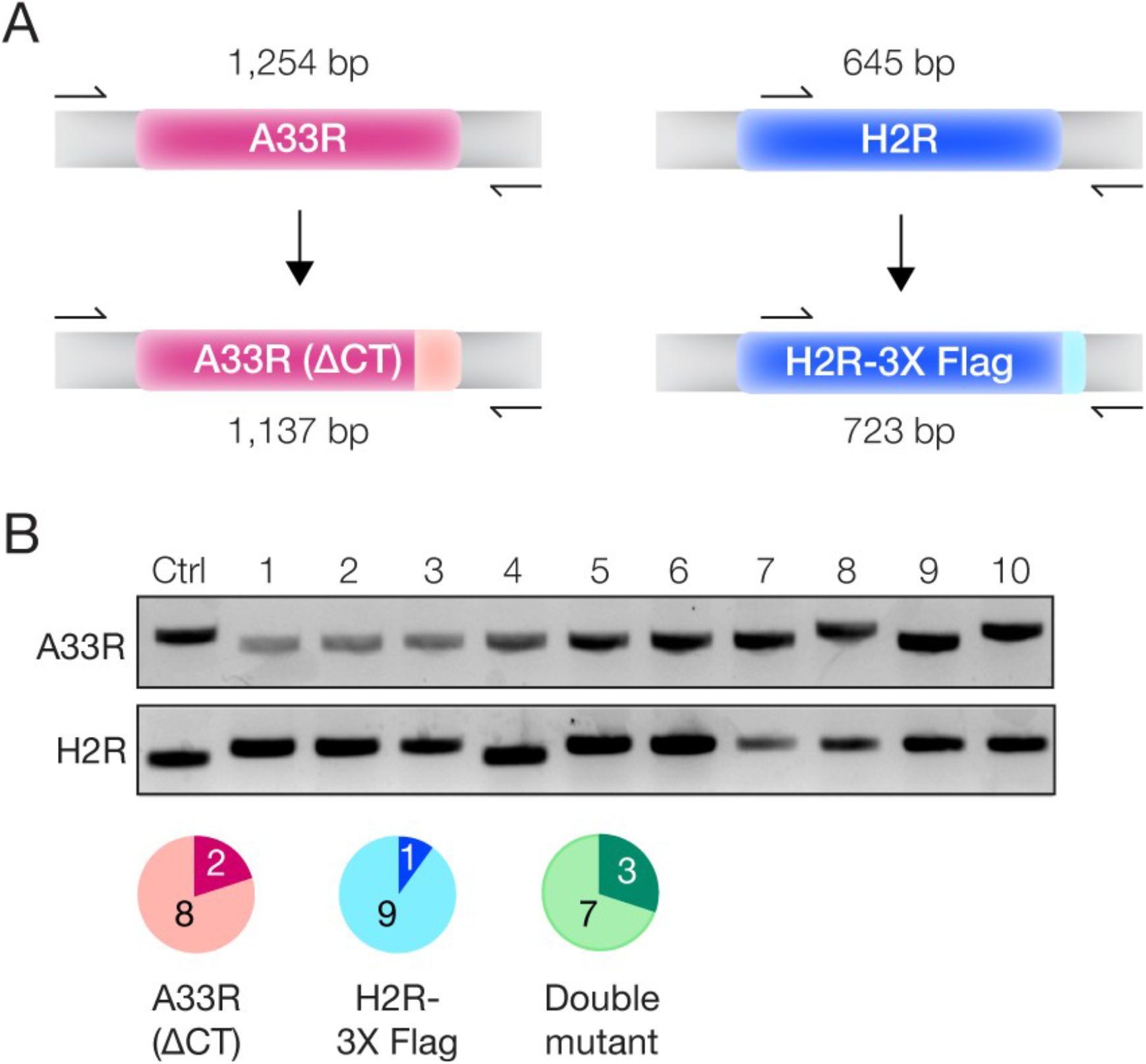
Simultaneous editing of two VACV genes with MAVERICC. (A) PCR amplification strategy. Primers (arrows) annealing outside the transfer amplicons were chosen to amplify the A33R or H2R loci in viral genomes, with differing PCR product lengths expected depending on whether or not the C–terminal truncation or FLAG tagging was successful. (B) PCR products generated by amplification of the A33R or H2R loci in 10 different progeny viral plaques or a WT control plaque (Ctrl). The pie charts show the number of plaques harboring the A33R mutation (pink), the H2R-3X FLAG tag insertion (blue), or both (green).

## Discussion

The genetic stability and manipulability of rVACVs, as well as their ease of propagation and broad host range, have made them popular as vectors for vaccine and immunotherapy applications. However, most methods for rVACV construction and rescue currently in use are slow, cumbersome, limited in versatility due to their need for markers or ‘landing pads’ in the viral genome, or dependent on customized reagents that pose a barrier to wider adoption. Here, we establish MAVERICC, a CRISPR/Cas9-based platform for rVACV engineering that overcomes many of these liabilities. We show that it affords the rapid and facile rescue of rVACVs modified at essentially any genomic locus without the need for markers or specially engineered parental viruses or cell lines. MAVERICC also sets the stage for the routine engineering of rVACVs at multiple, non-adjacent genomic loci in a single step and the construction of viral variant libraries at any genomic locus of interest.

The CRISPR/Cas9 system has been explored previously to engineer rVACVs. The first studies used CRISPR/Cas9 to cleave vDNA in cells to enhance the efficiency of viral rescue through homology-directed repair (HDR); however, they observed only incremental improvements over the traditional homologous recombination approach [11,12]. Thus, these workflows still required the use of selectable markers and multiple rounds of plaque purification. A more recent study demonstrated that Cas9 could rapidly and efficiently cleave vDNA in cells, but found that cleavage reduced viral replication and multiplication while having no stimulating effect on genome repair through homologous recombination [13]. This may be the case because (i) the cytoplasmic vDNA cannot be repaired by cellular HDR, which requires components that are localized to the nucleus; and (ii) efficient cleavage of vDNA by Cas9 soon after viral entry may inhibit VACV DNA replication, thereby also inhibiting homologous recombination, to which it is coupled. Indeed, whatever DNA replication initially occurs following Cas9 cleavage presumably arises from the subset of WT genomes that escape Cas9 action and is likely necessary to drive homologous recombination between the cleaved vDNA and transfer plasmid. As a consequence, only a small percentage of the resulting viral progeny pool is the desired rVACV, which must be enriched through multiple rounds of Cas9 selection against WT virus.

MAVERICC sidesteps these problems of incomplete Cas9 cleavage of intracellular vDNA in cells and the apparent trade-offs between vDNA cleavage, DNA replication, and homologous recombination in two ways. First, it temporally and spatially separates the processes of vDNA cleavage and homologous recombination. Performing vDNA cleavage with purified Cas9 *in vitro* affords a level of control, through the optimization of reactant concentrations, incubation time, and the use of multiple sgRNAs, that is not readily available to methods that cleave vDNA in infected cells. Moreover, any genomes that escape Cas9 cleavage in cells might be quickly sequestered into replication factories, creating a CRISPR-resistant pool of WT progeny virus. *In vitro* vDNA cleavage with CRISPR/Cas9 does not have to contend with such competing processes that could limit genome cleavage efficiency.

Second, like the restriction enzyme-mediated trimolecular recombination approach before it, MAVERICC decouples the source of the VACV genome (naked cDNA transfected into cells) from the helper virus (the avipoxvirus FWPV) that launches VACV DNA repair and replication. Because the VACV cDNA is non-infectious and FWPV is replication-defective in mammalian cells, their amounts could be independently optimized without concern for WT VACV background or helper virus-mediated cytopathic effects. Further, the recovery of rVACVs from both uncleaved and cleaved genomes in MAVERICC relies on FWPV, likely preventing the WT progeny generated from the few uncleaved vDNA molecules transfected into cells from outgrowing the recombinants. Our attempt to generate a virus bearing L5R-3X FLAG provides a striking case in point—despite the fact that this mutation appeared to be lethal, we observed little or no outgrowth of parental virus in the rescue experiment. In the CRISPR/Cas9-based approaches described above, by contrast, the parental VACV does double duty as both source of the rVACV genome and replication-competent helper virus, limiting the extent to which viral rescue can be optimized and yielding a pool of parental progeny as the unavoidable result.

MAVERICC enables applications that are not readily accessible via currently available approaches. First and foremost, it permits the engineering of any locus in the VACV genome in a straightforward manner without the need for selection/screening markers or specially prepared parental strains. Second, MAVERICC affords the parallel engineering of multiple loci and the simultaneous engineering of at least two loci with high efficiency, thereby creating unprecedented opportunities for genetic screens to investigate the basic biology of poxviruses and to identify viral variants with desirable properties for vector applications. In this regard, we note that we have successfully applied MAVERICC with all of the different gRNA pairs designed and tested so far (for this report and in other ongoing projects), indicating that the approach is robust and suggesting that strategies that require higher throughput are feasible. The increasing affordability of synthetic DNA technology should also facilitate such studies by simplifying the generation of repair amplicons. Third, MAVERICC extends the capabilities of the ‘classic’ trimolecular recombination approach for building rVACV libraries of individual genes. The latter relies on restriction enzyme digestion of vDNA *in vitro*, followed by helper virus-mediated rescue of recombinant viruses, and it affords libraries of up to 10^7^ unique viral variants for applications such as antibody discovery [17,26]. Our current work suggests that it should now be possible to construct such libraries of individual VACV genes as well.

## Acknowledgments

We thank Isabel Gutierrez, Estefania Valencia, and Laura Polanco for laboratory management and technical support. We thank Dr. Rohit Jangra, Robert Bortz and other members of the Chandran Laboratory for advice and suggestions and for comments on earlier versions of this manuscript. We thank Yakin Jaleta and Megan DeMouth for their independent tests of MAVERICC. We gratefully acknowledge Dr. Bernard Moss for helpful initial discussions and gifts of reagents and Dr. Stephen Laidlaw for advice on working with FWPV. This work was supported by National Institutes of Health (NIH) grant R01AI132633 (to K.C.). K.C. is a member of the scientific advisory boards of Integrum Scientific, LLC and the Pandemic Security Initiative of Celdara Medical, LLC.

## Materials and Methods

### Cells and viruses

BSC-40 cells were obtained from American Type Culture Collection (ATCC, CRL-2761) and were cultured in Dulbecco’s Modified Eagle Medium (DMEM, Gibco 11965-092) supplemented with 10% (vol/vol) fetal bovine serum (FBS, Atlanta biologicals S11150H), 1% (vol/vol) glutamine (Gibco 35050-061), and 1% (vol/vol) penicillin-streptomycin (Gibco 15140-122). The initial VACV, vNotI/tk, used in this study was a generous gift from Bernard Moss (NIH) [27]. This virus is a modified version of the Western Reserve strain that contains a unique *NotI* restriction site in the *thymidine kinase* (*tk*) gene. The vNotI/tk virus was propagated in BSC-40 cells and harvested by collection of the cell pellet after 2–3 days of growth and three freeze/thaw cycles to release the intracellular mature virions. It was then modified to contain either mCherry or eGFP reporters under control of the p11 VACV promoter in the *tk* locus. rVACV-mCherry and rVACV-eGFP were generated by transfecting a custom-synthesized plasmid containing 500 bases of left- and right-homology arms flanking the mCherry or eGFP reporter, followed by three rounds of plaque purification with selection for mCherry or eGFP expression.

Fowlpox virus (FWPV) was obtained from ATCC (VR-250) and propagated in chicken embryonic fibroblasts (CEFs) also obtained from ATCC (CRL-1590). CEFs were cultured in DMEM supplemented with 2.5% (vol/vol) FBS (Atlanta biologicals S11150H), 1% (vol/vol) glutamine (Gibco 35050-061), 1% (vol/vol) penicillin-streptomycin (Gibco 15140-122) and 1 mM HEPES (Gibco). CEFs were infected with FWPV when they were about 70% confluent and virus propagation was allowed to continue for 3 days or until cytopathic effects were evident throughout the cell culture. This FWPV strain initially did not grow well in these CEFs and therefore first had to be adapted to these cells through 9 serial passages. For each passage, a 15-cm plate of CEFs was grown to 70% confluence and 10% of the viral stock from the previous passage was added to the cells. The virus was grown for 3 days before cells were harvested by scraping and subjected to 3 freeze/thaw cycles to free intracellular virions.

### VACV DNA purification

VACV DNA was purified according to published methods [20,21]. First, 20 15-cm plates of BSC-40 cells were grown to confluence and infected with either rVACV-mCherry or rVACV-eGFP. After 3 days, cells were harvested by centrifugation and resuspended in 36 mL of TKE buffer (10 mM Tris-Cl, pH 8.0, 10mM KCl, 5 mM EDTA). Resuspended cells were kept on ice for 10 min with periodic gentle vortexing. After this time, 4 mL of 10% Triton X-100 and 100 µL of beta-Mercaptoethanol (BME) was added to the resuspension, followed by another 10-min incubation on ice with periodic gentle vortexing. The nuclei were then pelleted by centrifugation at 1,500 xg for 10 min before the supernatants were spun at 20,000 xg for 30 min to pellet the virus. The virus was then resuspended in 1.6 mL of cold TE buffer (10 mM Tris-Cl, pH 7.5, 1 mM EDTA) using a 21-gauge needle. After resuspension, the following were added in order with gentle mixing: 2.8 mL of 54% sucrose (w/w), 30 µL BME, 100 µL proteinase-K (10 mg/mL) and the solution was incubated on ice for 15 minutes. Then, 500 µL of 10% SDS was added and the solution was incubated overnight at 37°C. The following day, 1 mL of 4M NaCl was added with gentle mixing so as not to shear the vDNA. vDNA was then extracted three times with an equal volume of phenol:chloroform:isoamyl alcohol (25:24:1) by first gently mixing for several minutes, followed by a 3-min spin at 4,500 xg to separate the phases. The upper phase was saved after each spin and added to 550 µL of 3M sodium acetate, pH 7.0 after the last spin. Then, 12.5 mL of ice cold 100% ethanol was added with gentle mixing to this solution. The DNA was allowed to precipitate for at least 1 h at -80°C before being pelleted at 4,500 xg for 30 min. The pellet was washed once with 200 µL of 70% ethanol and spun again at 4,500 xg for 10 min. The pellet was allowed to air dry before being resuspended in 100 µL of TE buffer using pipette tips with the ends cut off to avoid DNA shearing. Several freeze/thaw cycles were conducted to ensure complete DNA resuspension before the concentration was determined with a Nanodrop spectrophotometer (ThermoFisher).

### Cleavage of vDNA with Cas9 and gRNAs

*Streptococcus pyogenes* Cas9-fused to a nuclear localization sequence (NLS) was obtained from Macrolab at UC Berkeley. sgRNAs were first assembled as DNA templates by PCR before *in vitro* transcription with the Megascript T7 RNA polymerase kit according to the manufacturer’s instructions. To assemble the DNA templates, forward and reverse DNA oligonucleotides (oligos) were ordered. The forward DNA oligos contained the T7 promoter (5’-TAATACGACTCACTATAGG-3’), followed by a 20-nucleotide sgRNA sequence (see below for details) and then a sequence to anneal to the 5’ end of the tracrRNA (5’-GTTTTAGAGCTAGAAATAGC-3’). Example forward oligo using the first mCherry guide (underlined) is listed below: 5’-TAATACGACTCACTATAGG<u>TATGCTATAAATGGTGAGCA</u>GTTTTAGAGCTAGAAATAGC-3’. A reverse DNA oligo designed to anneal to the 3’ end of the tracrRNA (5’-AAAAAAGCACCGACTCG-3’) was also ordered. PCR was then conducted using the forward and reverse DNA oligos to amplify each unique guide into a DNA template for *in vitro* transcription. For the PCR template, a plasmid containing the tracrRNA was used. PCR products were purified with the Qiagen PCR purification kit according to the manufacturer’s instructions before *in vitro* transcription.

To design sgRNAs, the CRISPOR program was used [19]. About 200 bp (NCBI Reference Sequence: NC_006998.1) surrounding the site of the desired insertion were submitted to the program. Potential sgRNAs were then manually mined to minimize off-target effects. sgRNAs that were in close proximity to the desired insertion site with minimal predicted off-targets were chosen. sgRNA sequences are listed in the supplemental Excel file.

For the Cas9 cleavage reaction, two gRNAs were used for each target gene. The Cas9, gRNAs and vDNA substrate were incubated in a 10:10:1 molar ratio with a final Cas9 concentration of 30 nM. The reaction was buffered in 1x New England BioLabs (NEB) buffer 3.1. The reaction was allowed to continue overnight at 37°C before being stopped by heat shock at 65°C for 5 min. To determine the Cas9 cleavage efficiency, 1.5 µg of the Cas9-treated vDNA was digested with either *HindIII*-HF or *XhoI* (NEB) for 3 h before being resolved on a large 0.6% agarose gel alongside an untreated control. DNA fragments were visualized with ethidium bromide staining. Throughout this process, cut pipette tips were used to avoid shearing the vDNA.

### Generation of transfer amplicons

To generate transfer amplicons for transfection, constructs were first custom-synthesized by either Epoch Life Science or Twist Biosciences and then amplified with primers designed to anneal to the ends of the synthesized sequences. For the eGFP rescue, the *eGFP* open-reading frame was placed under the control of the p11 promoter (5’-GAATTTCATTTTGTTTTTTTCTATGCTATAAATG-3’) and flanked on either side by 300 bp of homologous sequences at the *tk* locus. The primers to generate the transfer amplicon are listed in the supplemental Excel file. For *A33R* and *A34R*, the desired mutations were synthesized into a gene fragment flanked by about 500 bp of homologous sequences on each side. The specific mutations chosen were as follows; for *A33R*, the first 142 codons were unchanged, then a 12-nucleotide sequence (5’-TATCTAGCTCAT-3’) replaced the final 43 codons before the stop codon; for *A34R*, the 151st codon was changed from AAA to GAA. The primers to generate the transfer amplicons for *A33R* and *A34R* are listed in the supplemental Excel file. For *H2R* and *L5R*, a sequence encoding the 3X-FLAG tag was installed before the stop codon at the C– terminus of each gene and the coding sequence was flanked by 300 bp of homologous sequences on each side. The primers to generate the *H2R* and *L5R* are listed in the supplemental Excel file. After PCR of 30 cycles using Phusion polymerase, each amplicon was run on a 1% agarose gel and purified with the Qiagen gel extraction kit.

### Rescue of rVACVs

BSC-40 cells were seeded in wells of a 6-well plate at a density of ∼500,000 cells per well. The following day, each well was infected with 1.5 IU/cell of FWPV. Two hours later, 500 ng of cleaved vDNA was co-transfected with 150 ng of an appropriate transfer amplicon using Lipofectamine 3000 (Company) according to the manufacturer’s instructions. After overnight incubation, the media was replaced with DMEM-10% FBS and rVACV production was allowed to occur for 5–7 days. Cell supernatants and pellets were collected by scraping and viruses were released from the cells by three freeze/thaw cycles.

### TOPO cloning and sequencing

To estimate the efficiency of homologous recombination between the vDNA and transfer amplicon before proceeding to plaque purification, about 10% of the cell suspension mixed with the cell supernatant was harvested and vDNA was purified with the Qiagen blood mini prep kit according to manufacturer instructions. vDNA was then amplified by PCR with Phusion polymerase. Primers that anneal to the VACV genome outside the transfer amplicon were used to avoid re-amplifying the transfected construct. All “verification” primers used in this study are listed in the supplemental Excel file. After PCR, each amplicon was run on a 1% agarose gel and visualized with ethidium bromide staining.

We next sought to get a better estimation of what percent of the viral genomes had successfully incorporated the mutations for A33R and A34R. To assess this, Phusion polymerase and appropriate verification primers were used to amplify these loci with either 20 or 22 cycles, as determined in the supplemental figure. Three separate PCR reactions were performed for each sample, the products of each reaction were gel extracted and then reactions from the same sample were combined into one tube. The PCR pooled amplicons were then cloned into the pGEM-T plasmid using a TOPO cloning kit (Company) according to the manufacturer’s instructions. Individual bacterial colonies were then subjected to rolling-circle amplification and Sanger sequencing with gene-specific primers.

After purification and amplification of individual viral plaques, vDNA from each plaque was purified with a Qiagen blood mini prep kit. PCR was then conducted with matching verification primers, followed by gel extraction and Sanger sequencing of the PCR products with the same primers.

### Plaque assays

After initial virus rescue with the helper virus, individual plaques were grown to assess the efficiency of rVACV formation. After freeze/thawing, a 10-fold dilution series for each rescue was set up in a 6-well plate with a starting dilution of 10^−3^. One hour after infection, the media was exchanged with a 0.5% methylcellulose overlay and incubated for 2-3 days. Plaques were then randomly selected and the virus was expanded by growing on BSC-40 cells in a 6-well dish. For the assay to determine the size of the A33R plaques, the plaques were grown for 4 days under a 0.5% methylcellulose overlay and then stained with 0.1% crystal violet.

### Western blotting

BSC-40 cells were seeded into a 6-well plate and infected with virus. After two days, cells were collected by scraping and the cell pellet was lysed in 1% SDS. Proteins in 20 µg of each lysate were then resolved on a 4–20% polyacrylamide gradient gel and transferred to a PVDF membrane. Membranes were blocked in 4% milk and stained with anti-FLAG (1:2000 dilution) and anti-mouse-HRP (1:5000 dilution) primary and secondary antibodies. Protein bands were detected via chemiluminescence using a Bio-Rad ChemiDoc Touch imager.

### Immunofluorescence

BSC-40 cells were seeded onto a coverslip in a 12-well plate and infected with virus. The following day, cells were harvested by fixation for 15 min with 4% paraformaldehyde. Cells were washed 2x with PBS and then permeabilized with 0.1% Triton X-100 in PBS for 10 min. Cells were washed again with PBS and blocked with 3% BSA in PBS for 10 minutes. A primary anti-FLAG antibody was diluted 1:500 in 3% BSA/PBS and incubated with the coverslip for 1 h. Cells were then washed 5 times with PBS and incubated for 1 h with secondary antibody, which had been diluted 1:1000 in 3% BSA/PBS. Cells were then washed 4 times with PBS, with the penultimate wash containing Hoechst dye at a 1:20,000 dilution. Coverslips were mounted onto slides with Prolong Gold (Company) as a mounting reagent and sealed with nail polish. Images were taken at either 40x or 63x magnification using a Zeiss Axio Observer inverted microscope.

---

**Supplemental figure 1.**
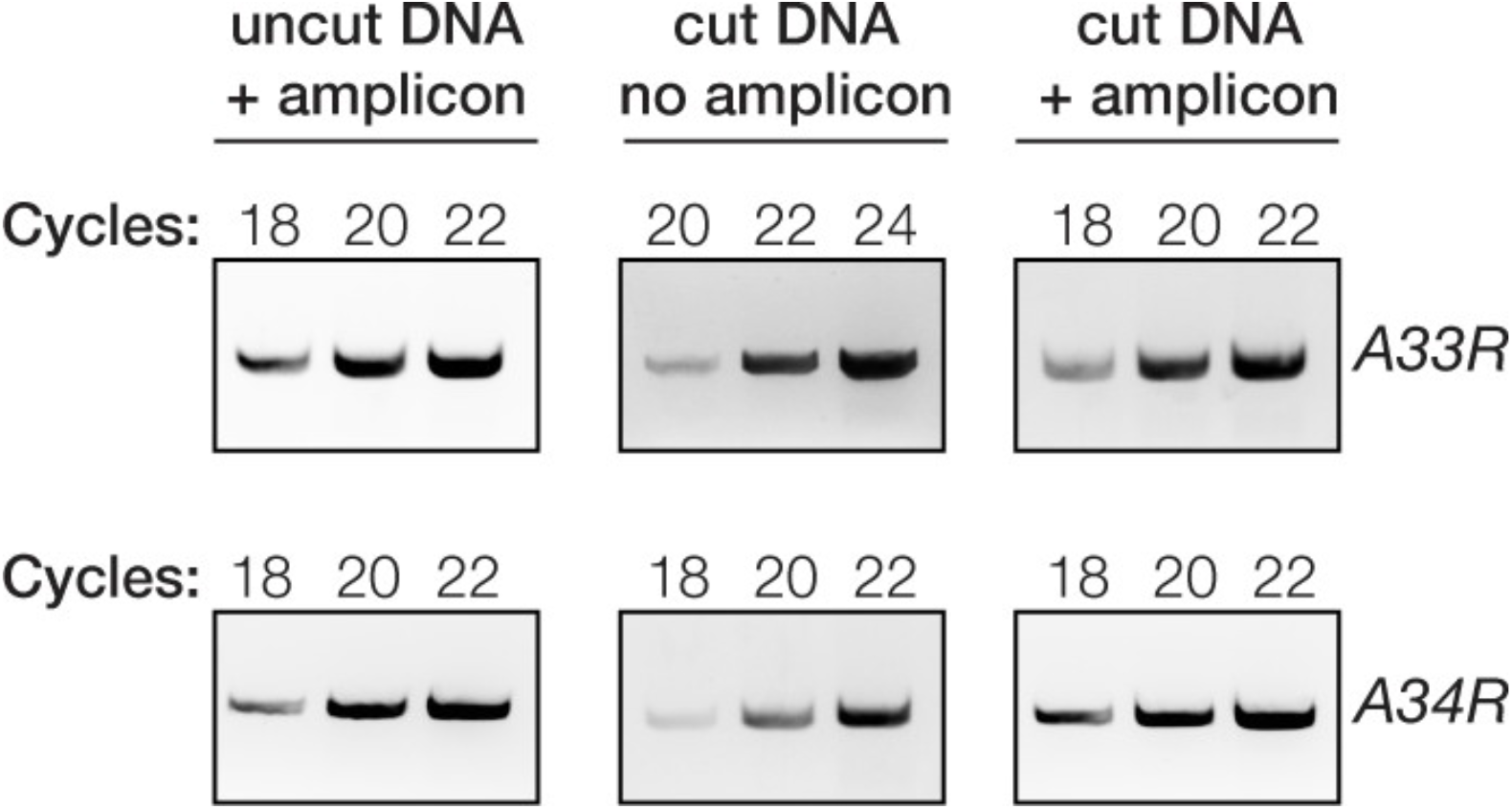
Determination of the number of PCR cycles to use for TOPO cloning and sequencing in Fig 2D. vDNA was prepared by miniprep from MAVERICC A33R or A34R rescue reactions after 5 days. Three different numbers of PCR cycles were then performed for each sample and analyzed by agarose gel electrophoresis. For each sample, the middle number of cycles from each gel was chosen to proceed to clone the PCR amplicon into the pGEM-T vector.

